# Machine learning-based prediction of dynamic height heterosis with pathway biomarkers in rice

**DOI:** 10.1101/2024.11.09.622823

**Authors:** Zhiwu Dan, Yunping Chen, Wenchao Huang

**Affiliations:** State Key Laboratory of Hybrid Rice, Key Laboratory for Research and Utilization of Heterosis in Indica Rice, the Ministry of Agriculture, The Yangtze River Valley Hybrid Rice Collaboration & Innovation Center, College of Life Sciences, Wuhan University, Wuhan 430072, China

**Keywords:** dynamic heterosis, pathway biomarker, metabolomics, plant height, rice

## Abstract

The development of robust biomarkers enables accurate prediction of complex phenotypes. However, the dynamic nature of biomarkers is often underestimated since their quantitative changes during development are directly connected to phenotypic transformations, influencing both crop agronomic traits and human diseases. Here, we performed network analysis of untargeted metabolite profiles to investigate height heterosis in rice, which is dynamic that varies during development and is a key determinant of yield heterosis. We found that the levels of pyruvaldehyde were predictive of height heterosis specific at the seedling stage, while 4-hydroxycinnamic acid positively correlated with height heterosis across four developmental stages. We identified metabolic pathways associated with height heterosis and found that metabolomic changes during the elongation stage had a greater impact than those in other stages. Finally, 11 heterosis-associated pathways were developed into metabolomic biomarkers through random forest analysis, successfully predicting height heterosis in an independent population under different growth conditions. This study elucidates the metabolomic landscape of dynamic height heterosis in rice and develops pathway biomarkers for complex phenotypes, demonstrating robustness across diverse populations, environments, and developmental stages.

## Introduction

Complex phenotypes are mainly determined by genotypes, environmental factors, and their interactions (Hunter, 2005; Crossa et al., 2017; Scheres and van der Putten, 2017). The influence of genetic and environmental factors on phenotypic changes has been extensively studied in both crop agronomic traits and human diseases (Fave et al., 2018; Li et al., 2018; de Los Campos et al., 2020). To accurately characterize the states of complex phenotypes, various types of biomarkers have been developed, and predictive models have been constructed to facilitate accurate predictions across populations and environments (Riedelsheimer et al., 2012; Sarkizova et al., 2020; Tong et al., 2020; Zhao et al., 2021). However, the dynamic nature of the processes shaping complex phenotypes is often underestimated (Lesterhuis et al., 2017; Miao et al., 2020). Quantitative changes in specific molecules during development have been shown to directly linked to phenotypic transformations, including plant agronomic traits (Obermeyer et al., 2013; Ko et al., 2016; Varoquaux et al., 2019; Zhou et al., 2019a) and risks for human diseases such as diabetes or cancers (Taylor et al., 2009; Price et al., 2017). Consequently, static biomarkers—identified from tissues at a single developmental stage—often exhibit limited predictive power for new-stage phenotypes in genetically distant individuals, partly due to molecular heterogeneity across developmental stages and genotypes (Menche et al., 2017; Price et al., 2017). Furthermore, the question of at which stages quantitative changes in biomarkers contribute more to the phenotypic outcome than at others remains largely unexplored. Thus, investigating the molecular mechanisms of complex phenotypes in a time series manner, followed by the exploration and validation of dynamic biomarkers, offers a viable approach for advancing precision breeding and precision medicine.

Plant height is a key determinant of crop grain yield (Wang et al., 2023). Although plant height is not a component trait of grain yield, it significantly contributes to yield through its associations with grain number and flowering time. In rice, the close relationship between plant height and yield has been attributed to multiple polygenic genes, such as *Ghd7* and *DTH* (Xue et al., 2008; Gao et al., 2014). In a partial correlation analysis comparing the contributions of four yield components and plant height to yield heterosis in rice, heterosis for plant height (*R* = 0.22) had a higher correlation coefficient than that of grain weight (*R* = 0.16) (Dan et al., 2021). Notably, *Ghd7* plays a central role in yield heterosis of the elite hybrid rice Shanyou63 (Wang et al., 2022). However, plant height is a dynamic trait that evolves from seedling to maturity, yet most studies primarily focus on the maturation stage. Given the substantial increase in crop grain yield achieved through heterosis, investigating the molecular mechanisms underlying height heterosis has become essential for further enhancing yield potential.

In this study, we measured plant height in rice diallel crosses across four developmental stages: seedling, elongation, flowering, and maturation. We observed a significant shift in height heterosis between the elongation and flowering stages. Subsequently, we identified heterosis-associated analytes at each stage and examined their relationships with height heterosis. We also detected metabolic pathways involved in dynamic height heterosis and discovered that the overlapping pathways—primarily in amino acid and carbohydrate metabolism—connected different developmental stages. Notably, we observed pronounced metabolomic changes at the elongation stage. Using random forest, we developed pathway biomarkers and validated them with test crosses. Our findings provide insights into the metabolomic mechanisms of dynamic height heterosis and demonstrate the potential of pathway biomarkers in predicting complex phenotypes across developmental stages.

## Results

### Dynamic heterosis for plant height of diallel crosses in rice

To investigate the molecular mechanisms underlying dynamic heterosis in rice, we selected 18 representative rice inbred lines, which consist of both *indica* and *japonica* varieties with varying plant height (Dan et al., 2021; Dan et al., 2024a; Dan et al., 2024b), as parental lines. Using a complete diallel design, we generated a hybrid population and measured plant height of the population (including both inbred lines and F_1_ hybrids) at four developmental stages: seedling stage (SS), elongation stage (ES), flowering stage (FS), and maturation stage (MS). Plant height data of 18 inbred lines and a total of 287 F_1_ hybrids were successfully collected, and better-parent heterosis for plant height (BPH_PH) were calculated for the hybrid population.

Height heterosis varied among individuals across the four stages (Fig. 1A and Supplementary Table S1). When analyzed collectively, the F_1_ hybrids exhibited significantly stronger heterosis at later stages compared to earlier stages (Fig. 1B). Additionally, we classified the F_1_ hybrids into high- and low-BPH groups based on the 75th and 25th percentiles of heterosis at each stage for subsequently comparative analysis. Despite stage-specific variations, heterosis was positively correlated across stages, with correlation coefficients decreasing as the interval between stages increased (Fig. 1C).

**Figure 1.**
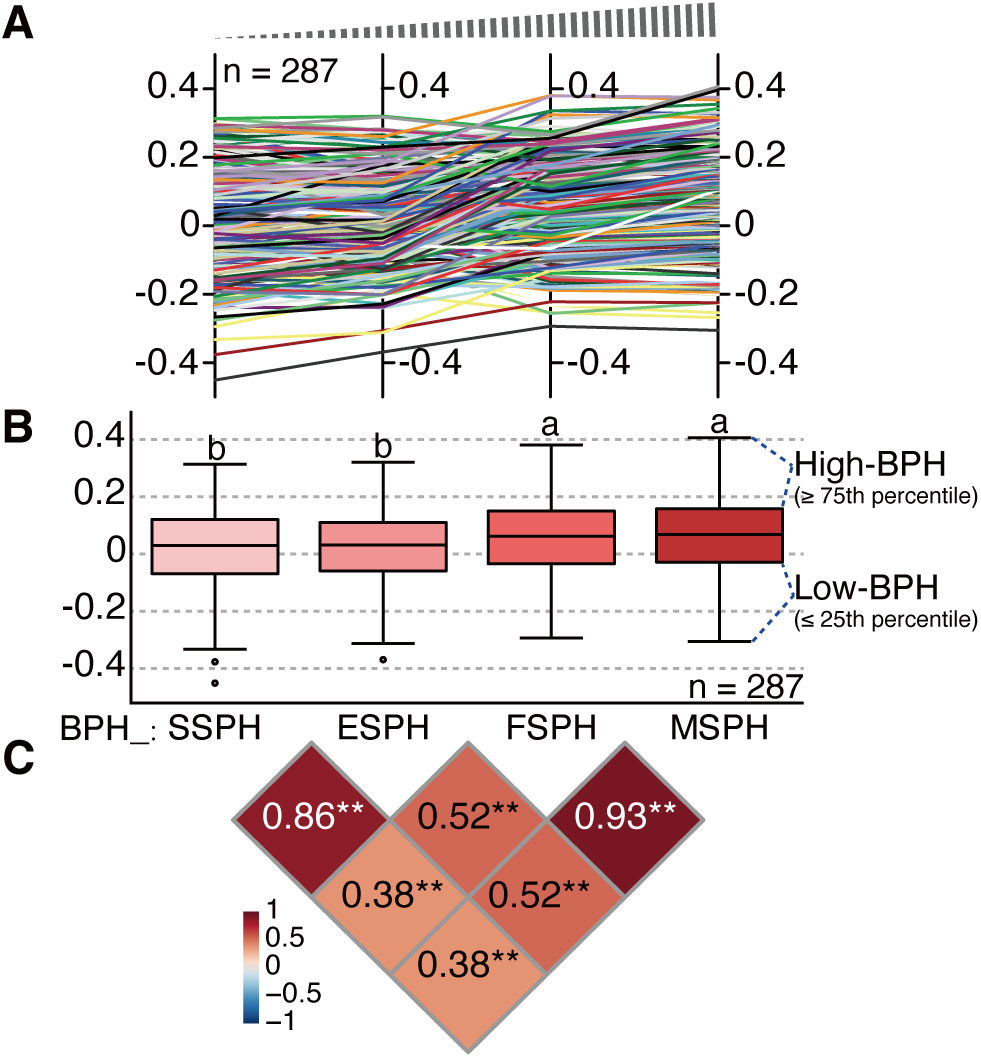
Dynamic heterosis for plant height of the diallel crosses. **(A-B)** Better-parent heterosis for plant height of the F_1_ hybrids from the diallel cross population at the individual (**A**) and populational levels (**B**). The F_1_ hybrids with better-parent heterosis (BPH) ≤ 25th and ≥ 75th percentiles at each stage were grouped as low- and high-BPH hybrids, respectively. For box plots in (**B**), the middle line is the median. The lower and upper boundaries of the boxes are the first and third quantiles, respectively. The whiskers represent the 1.5 times interquartile range, and the circles represent the outliers. Different letters above the boxes in (**B**) indicate significant differences between groups (one-way ANOVA with the least significant difference test for post hoc multiple comparisons, *P* < 0.05). **(C)** Pearson correlations among height heterosis across the four developmental stages. Abbreviations: SSPH, seedling stage plant height; ESPH, elongation stage plant height; FSPH, flowering stage plant height; MSPH, maturation stage plant height. **, significant at the 0.01 level.

### Metabolic analytes associated with dynamic height heterosis

To explore the metabolic underling of dynamic heterosis, we integrated untargeted metabolite profiles with height heterosis across the four stages. We first obtained metabolite profiles from young seedlings of the parental inbred lines to calculate metabolite levels for F_1_ hybrids. Since additive effect, namely at middle parental levels, is among the predominant inheritance patterns in F_1_ hybrids (Zhou et al., 2019a; Liu et al., 2021; Dan et al., 2024a; Dan et al., 2024b), we averaged the parental metabolite levels to represent the hybrid metabolome as previously described (Dan et al., 2019; Dan et al., 2020). Subsequently, we employed a partial least square-based strategy to connect the calculated hybrid metabolome and dynamic heterosis (Wold, 1975; Dan et al., 2021). Predictive models for height heterosis of each stage were built and these models displayed best performance with four latent factors (Fig. 2A). The top *r* values of these predictive models ranged from 0.68 to 0.77, indicating high accuracy of the additive effect for representing metabolic levels of F_1_ hybrids.

**Figure 2.**
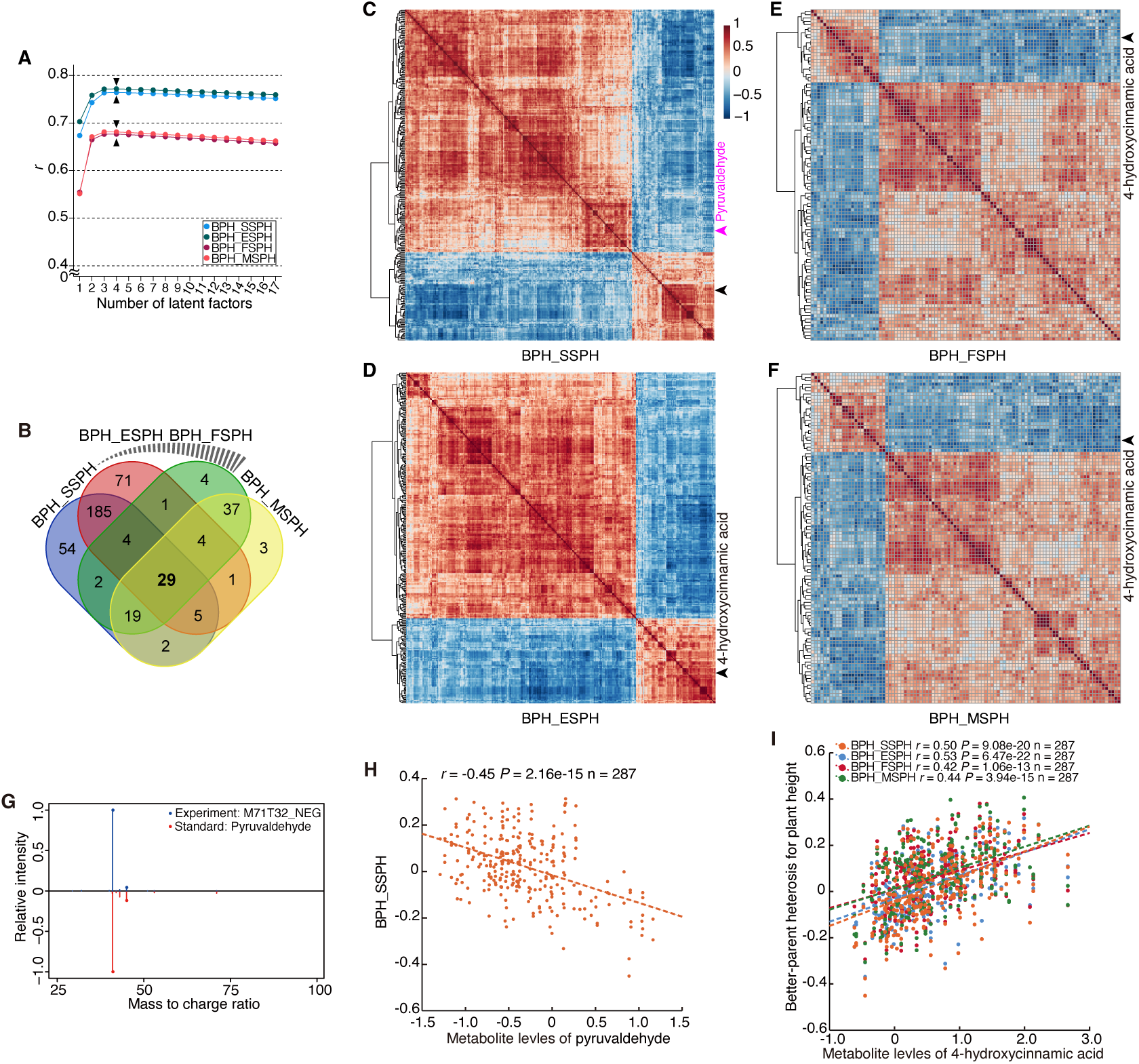
Heterosis-associated analytes for dynamic height heterosis. **(A)** Changes in *r* values with varied number of latent factors. Additive effect was adopted to calculate metabolite levels of F_1_ hybrids. Predictive models were built for height heterosis at four stages with the partial least square method. **(B)** Venn diagram of heterosis-associated analytes across four developmental stages. **(C-F)** Heatmaps of correlations among the heterosis-associated analytes for plant height at the seedling (**C**), elongation (**D**), flowering (**E**), and maturation (**F**) stages, respectively. Pearson correlation was performed among analytes. The locations of analytes annotated as 4-hydroxycinnamic acid and pyruvaldehyde are marked with black (at four stages) and purple (only the seedling stage) arrows, respectively. **(G)** MS/MS spectra of M71T32_NEG and the pyruvaldehyde standard. **(H)** Correlations between metabolite levels of pyruvaldehyde and height heterosis at the seedling stage. **(L)** Correlations between metabolite levels of 4-hydroxycinnamic acid and height heterosis at four developmental stages.

Contributions of each analyte in predictive models were assesses by average values of variable importance in projection, and the top contributed analytes were identified for heterosis at each stage. The final numbers of heterosis-associated analytes for the four stages were 300, 300, 100, and 100, respectively, suggesting the involvement of more metabolic pathways with early-stage heterosis (Fig. 2B).

We performed correlation analysis on these heterosis-associated analytes to investigate their correlation patterns. Both positive and negative correlations were found and most of these analytes had positive correlations with other analytes across the four stages (Fig. 2C-F). Notably, an accurately annotated metabolite pyruvaldehyde (peak tag: M71T32_NEG; Fig. 2G), which is involved in pyruvate metabolism and propanoate metabolism, showed a significant negative correlation with height heterosis at the seedling stage (Fig. 2H). In contrast, 4-hydroxycinnamic acid, which has been proven to be linked to plant height (Li et al., 2015; Dan et al., 2021), was consistently identified as a heterosis-associated analyte across all four stages (Fig. 2C-F). The levels of 4-hydroxycinnamic acid showed significant positive correlations with plant height heterosis throughout the stages, with the later stages exhibiting lower correlation coefficients compared to the earlier ones (Fig. 2J).

### Metabolic pathways underlying dynamic height heterosis

Due to limited annotation information of the heterosis-associated analytes, it was challenging to classify them into metabolic pathways. We thus adopted dysregulated network analysis (Shen et al., 2019), which involves pathway enrichment analysis using a hypergeometric test on the differential metabolic peaks, to identify heterosis-associated metabolic pathways. By treating the low- and high-BPH hybrids as control and case groups, respectively, we obtained significantly enriched pathways (heterosis-associated pathways) for each stage. Consistent with the trend in numbers of heterosis-associated analytes, more pathways were implicated in height heterosis at early stages (Fig. 3A-D). Through metabolic levels of these pathways, namely the average levels of all metabolites in a pathway (Shen et al., 2019), we performed correlation analyses and observed similar correlation patterns of these pathways to those of heterosis-associated analytes (Fig. 3A-C).

**Figure 3.**
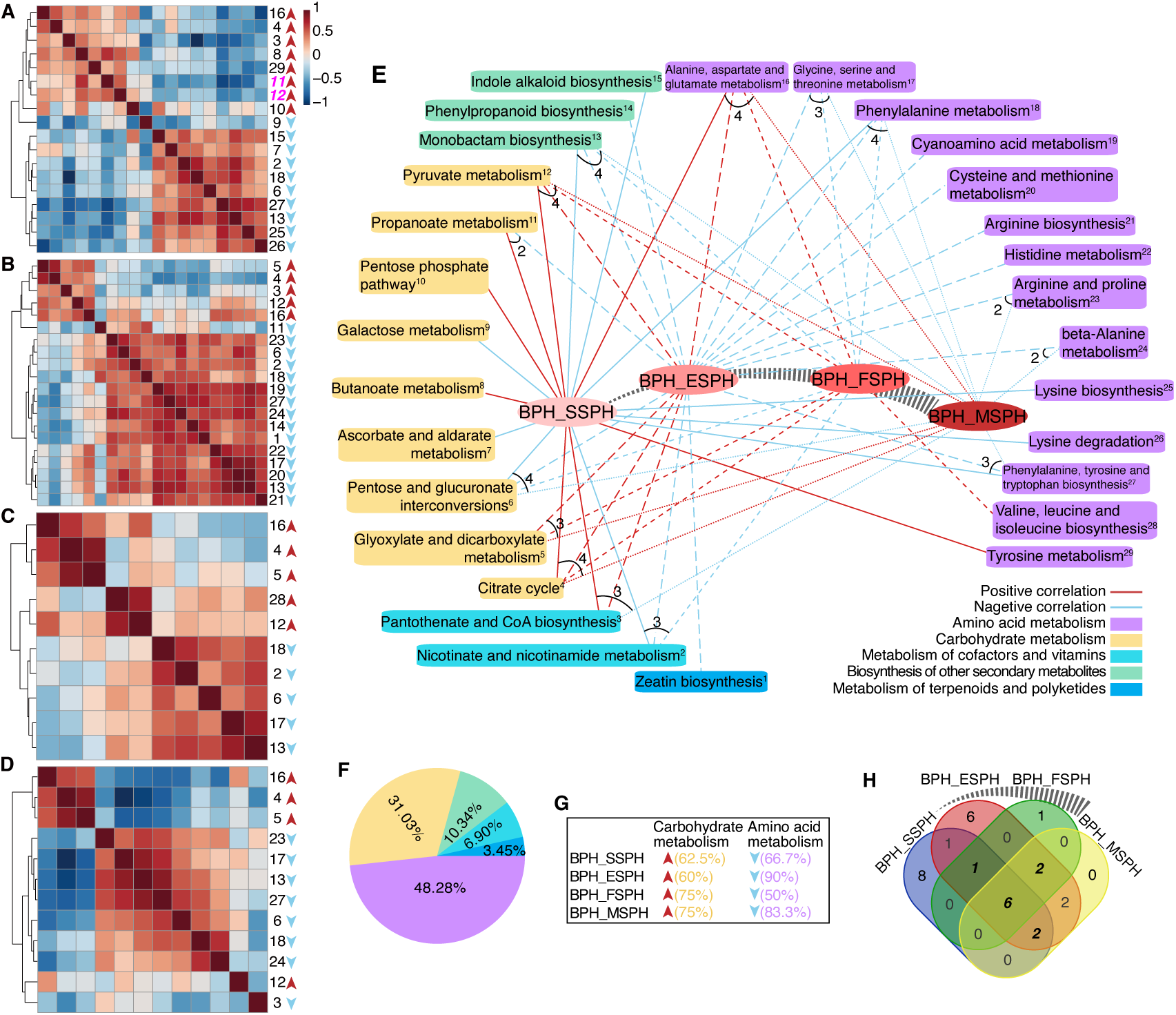
Heterosis-associated pathways underlying dynamic height heterosis. **(A-D)** Heatmaps of correlations among the heterosis-associated pathways for height heterosis at the seedling (**A**), elongation (**B**), flowering (**C**), and maturation (**D**) stages, respectively. The metabolite levels of the pathways for height heterosis were obtained through dysregulated network analysis on the high- and low-BPH hybrids. **(E)** The metabolomic landscape of dynamic height heterosis. Pearson correlations were performed between height heterosis and metabolite levels of the heterosis-associated pathways. **(F)** The pathways were from five groups with different percentages. **(G)** Percentages of the pathways in carbohydrate metabolism that had high metabolite levels in the high-BPH hybrids (upward red arrow) and those of the pathways in amino acid metabolism that had low metabolite levels in the high-BPH hybrids (downward light blue arrow). **(H)** Venn diagram of the heterosis-associated pathways for height heterosis across four developmental stages. The 11 metabolic pathways that were significantly enriched at most stages were among the heterosis-associated pathways for plant height at the elongation stage.

Among these pathways, pyruvate metabolism and propanoate metabolism were associated with height heterosis, while only pyruvate metabolism was also involved in heterosis at the later stages. Pyruvate metabolism exhibited a positive correlation with height across all four stages, whereas propanoate metabolism showed a positive correlation at the seedling stage but a negative correlation at elongation. These findings suggested that the relationships between metabolic pathways and heterosis might differ from those of individual analytes, underscoring the importance of exploring height heterosis at the pathway level.

By interconnecting overlapping pathway across stages, we obtained the metabolomic landscape of dynamic height heterosis (Fig. 3E). A total of 29 metabolic pathways were detected, with three, five, and six metabolic pathways involved in height heterosis at two, three, and all four stages, respectively. The majority of these heterosis-associated pathways belonged to amino acid (48.28%) and carbohydrate (31.03%) metabolism (Fig. 3F). Notably, the pathways in carbohydrate metabolism (60-75%) consistently exhibited high metabolite levels in high-BPH hybrids, while those in amino acid metabolism (50-90%) displayed lower levels (Fig. 3G). This pattern underscored the dominant roles of carbohydrate and amino acid metabolism in height heterosis, which complements the changing trends observed for metabolic pathways related to heterosis of yield and its components (Dan et al., 2020; Dan et al., 2021).

Furthermore, the numbers of heterosis-associated pathways were higher for the earlier stages (8 and 6 pathways) than the later stages (1 and 0 pathway; Fig. 3H), suggesting the changes in heterosis-associated pathways drove height heterosis variation. However, the contributions of these heterosis-associated pathways at each stage to height heterosis remained unclear.

### Metabolomic changes at the elongation stage play critical roles in height heterosis

According to the metabolomic landscape of height heterosis, six pathways were significantly enriched across all four stages. Three of these pathways (citrate cycle, pentose and glucuronate interconversions, and pyruvate metabolism) were classified under carbohydrate metabolism, while two pathways were from amino acid metabolism (phenylalanine metabolism and alanine, aspartate and glutamate metabolism). The remaining pathway was monobactam biosynthesis. To assess metabolomic variations linked to dynamic heterosis, we compared metabolite levels of these pathways across stages (Fig. 4A-F). Significant differences in metabolite levels of the six pathways were observed in either the high- or low-BPH hybrids across the four stages. Nearly all pathways exhibited metabolomic differences in at least one stage when compared to the others. In high-BPH hybrids, the citrate cycle showed an initial increase at elongation followed by a decline in later stages (Fig. 4A), whereas the opposite trend occurred in low-BPH hybrids. Similar changing patterns were observed for the other two pathways from carbohydrate metabolism (Fig. 4B-C). In contrast, the two pathways from amino acid metabolism and monobactam biosynthesis displayed fewer changes in metabolite levels in low-BPH hybrids (Fig. 4D-F). Notably, metabolite levels frequently shifted significantly at the elongation stage in both high- and low-BPH hybrids, underscoring its critical role in shaping height heterosis.

**Figure 4.**
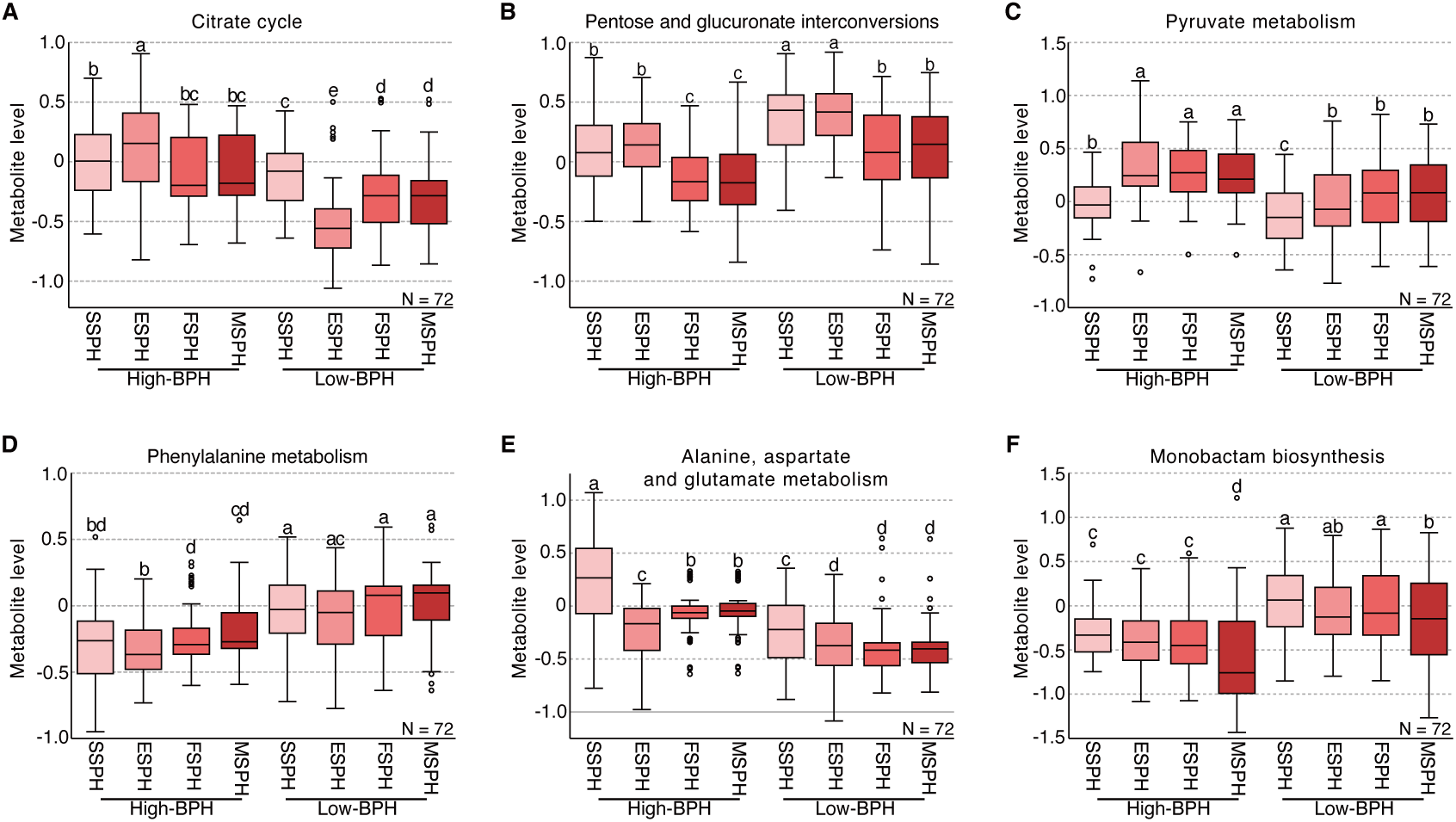
Metabolite levels of six metabolic pathways in the high- and low-BPH hybrids. Metabolite levels of six metabolic pathways, including the citrate cycle (**A**), pentose and glucuronate interconversions (**B**), pyruvate metabolism (**C**), phenylalanine metabolism (**D**), alanine, aspartate and glutamate metabolism (**E**), and monobactam biosynthesis (**F**), were compared to investigate changing patterns across stages. For box plots, the middle line is the median. The lower and upper boundaries of the boxes are the first and third quantiles, respectively. The whiskers represent the 1.5 times interquartile range and the circles represent the outliers. Different letters above the boxes indicate significant differences between groups (one-way ANOVA with the least significant difference test for post hoc multiple comparisons, *P* < 0.05).

### Pathway biomarkers for predicting dynamic height heterosis

To explore biomarkers for the dynamic height heterosis, we first performed univariate and multivariate receiver operating characteristic (ROC) curve analyses on the 18 heterosis-associated metabolic pathways for height heterosis at the seedling stage (Supplementary Fig. S1). A single predictor achieved an area under the curve (AUC) of 0.885, increasing to 0.933 with 100 features. Analysis of the six heterosis-associated pathways across all stages revealed no substantial increase in AUC values or predictive accuracies (uni-ROC: AUC = 0.887, multi-ROC: AUC = 0.931, predictive accuracy = 0.882 with 21 features; Supplementary Fig. S1).

We then evaluated the 11 heterosis-associated pathways that were for three or four stages, which were primarily pathways in amino acid and carbohydrate metabolism, as predictive features (Fig. 5A). A single predictor yielded an AUC was 0.883 (Fig. 5B), while the optimal predictive model (10 features) achieved an AUC of 0.929 and predictive accuracy of 0.873 (Fig. 5C-D). Unexpectedly, despite being significantly enriched across all four stages, the citrate cycle was not among the top-ranked ten predictors (Fig. 5E). Further biomarker analysis demonstrated high predictive accuracies for height heterosis at the three later stages: elongation, flowering, and maturation (Supplementary Fig. S2). These results showed that the development of biomarkers from the metabolic pathways associated with heterosis across most developmental stages could be an effective strategy, taking into account predictive accuracy, feature number, and feature ranking.

**Figure 5.**
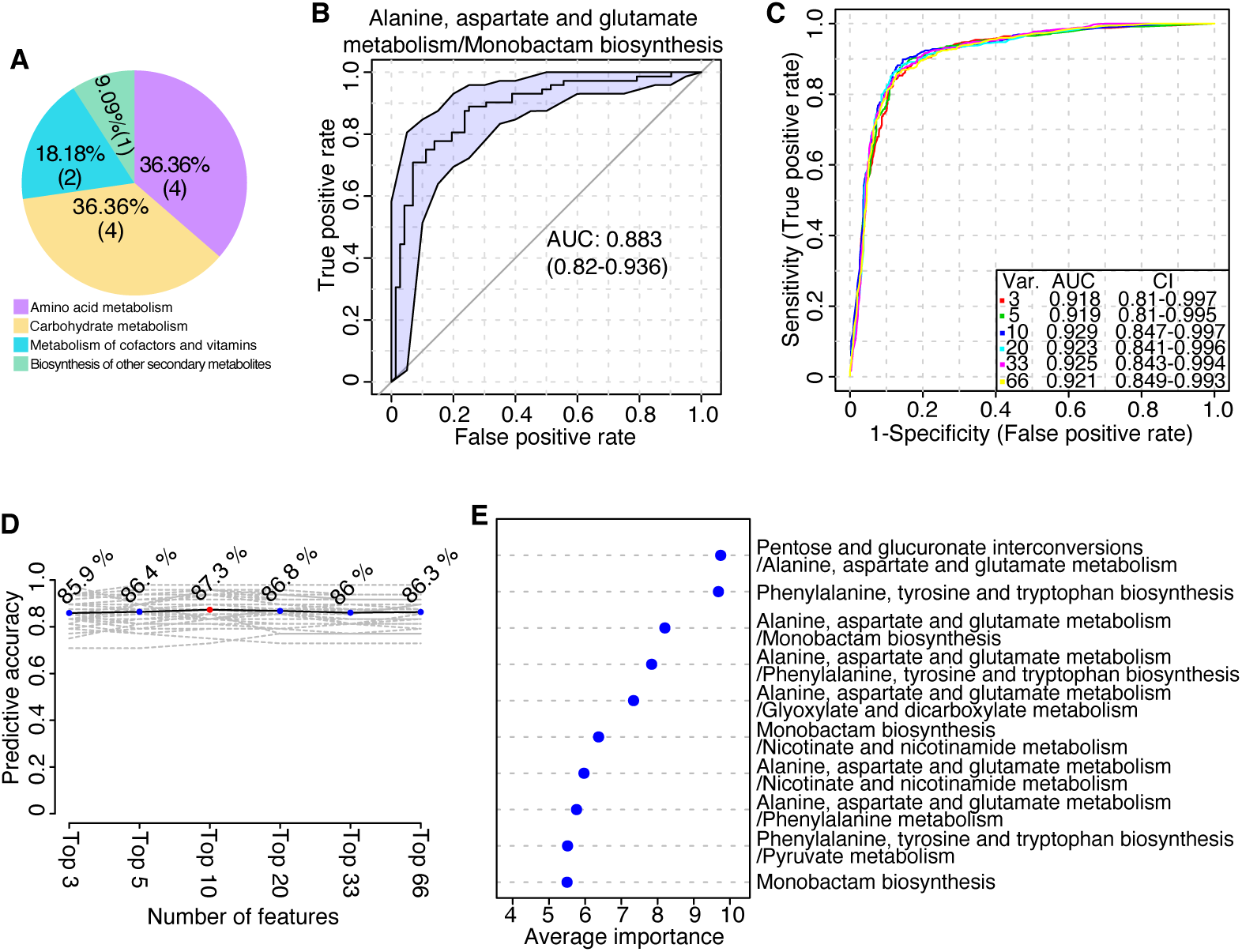
The development of pathway biomarkers for dynamic height heterosis. **(A)** Categories of the 11 metabolic pathways that were significantly enriched at most (three and four) developmental stages. **(B)** Univariate receiver operating characteristic (ROC) curve analysis of height heterosis at the seedling stage of the diallel crosses. The metabolic ratios of alanine, aspartate and glutamate metabolism and monobactam biosynthesis were calculated and used as predictors. AUC, area under the curve. **(C-E)** Multivariate ROC curve analysis of height heterosis at the seedling stage of the diallel crosses. The changes in the AUC and predictive accuracies with different numbers of variables (VAR.) or features are shown in (**C**) and (**D**), respectively. The average importance of the top ten features is listed in (**E**). CI, confidence interval.

### Validation of pathway biomarkers with test crosses

To validate predictive capability of these pathway biomarkers, we utilized height heterosis of a group of Honglian-type test crosses, which exhibited distant genetic relationships and were cultivated under different field conditions than the diallel crosses (Dan et al., 2020). The test crosses exhibited a narrower range of height heterosis due to limited genetic diversity of their male parents, which were mainly recombinant inbred lines (Fig. 6A). Meanwhile, the height heterosis (at the maturation stage) of the test crosses was comparable to those of the diallel ones. After categorizing the test crosses into low- and high-BPH groups based on the 25th and 75th percentiles of height heterosis, we obtained metabolite levels for four pathways through dysregulated network analysis (Supplementary Table S2), three of which overlapped with the 11 previously identified pathways. Using the metabolite levels of nicotinate and nicotinamide metabolism as a single predictor yielded an AUC of 0.868 (Fig. 6B), while AUC values ranging from 0.8 to 0.834 were achieved with 2-6 variables (Supplementary Fig. S3).

**Figure 6.**
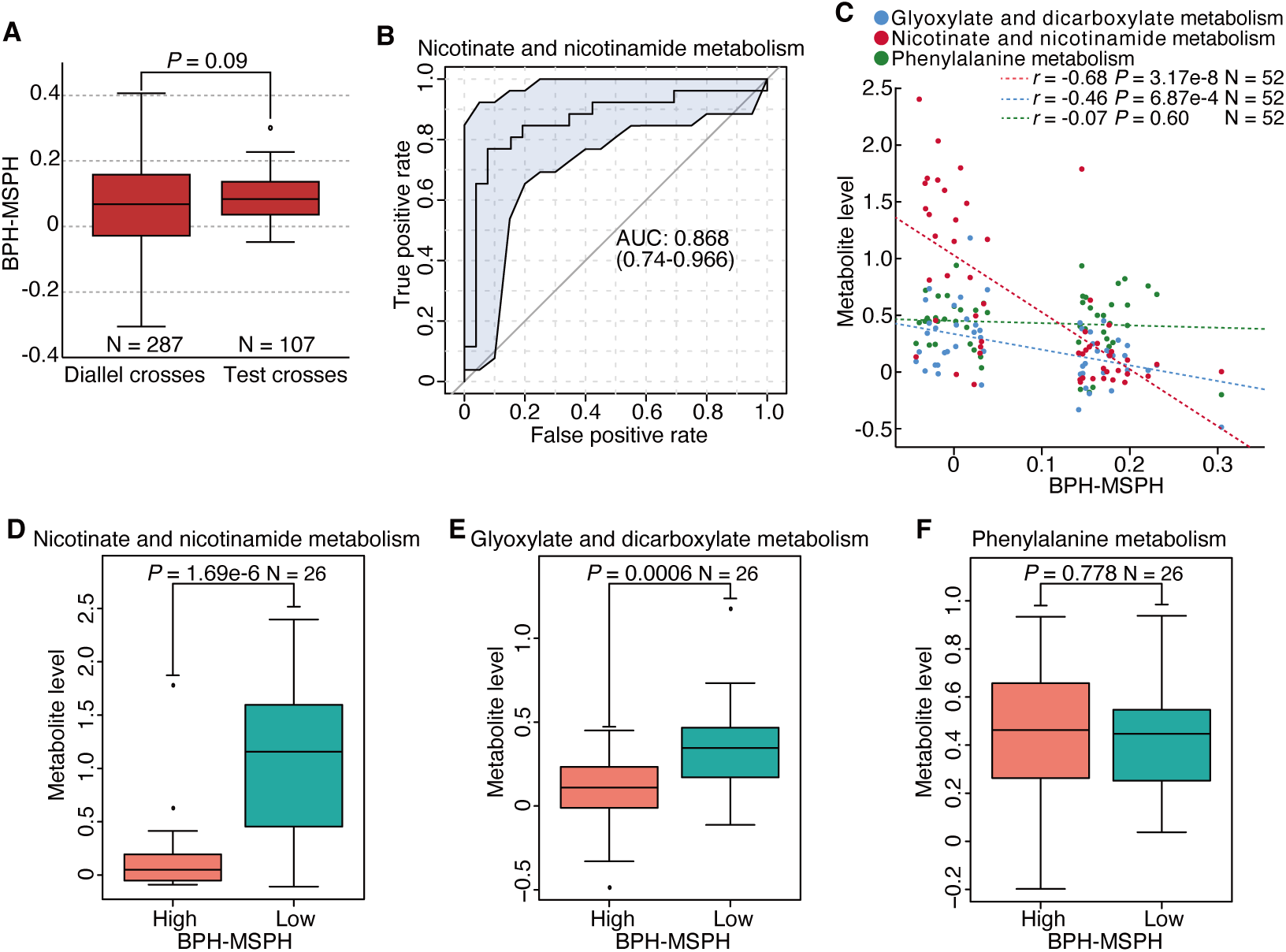
Validation of pathway biomarkers in test crosses. **(A)** Box plots of plant height heterosis at the maturation stage of the diallel and test crosses. **(B)** Univariate ROC curve analysis of height heterosis at the maturation stage of the test crosses. The test crosses were divided into low- and high-BPH hybrids based on the 25th and 75th percentiles, respectively. The metabolite levels of nicotinate and nicotinamide metabolism were identified as predictors. **(C)** Pearson correlations between height heterosis at the maturation stage of the test crosses and metabolite levels of three metabolic pathways. **(D-F)** Comparisons of the metabolite levels of three pathways between the high- and low-BPH hybrids from the test crosses. The metabolite levels of nicotinate and nicotinamide metabolism (**D**), glyoxylate and dicarboxylate metabolism (**E**), and phenylalanine metabolism (**F**) were compared. In (**A**) and (**D**-**F**), the data were analyzed by two-tailed independent samples *t*-test. For the box plots in (**A**) and (**D**-**F**), the middle line is the median. The lower and upper boundaries of the boxes are the first and third quantiles, respectively. The whiskers represent the 1.5 times interquartile range, and the circles represent the outliers.

Height heterosis was negatively correlated with the metabolite levels of nicotinate and nicotinamide metabolism, as well as glyoxylate and dicarboxylate metabolism (Fig. 6C). But no correlation was found between height heterosis and phenylalanine metabolism. Additionally, the metabolite levels of nicotinate and nicotinamide metabolism were significantly lower in the high-BPH hybrids compared to low-BPH hybrids (Fig. 6D). The levels of glyoxylate and dicarboxylate metabolism were also lower in the high-BPH hybrids (Fig. 6E), which contrasted with the result shown in Fig. 3E. Phenylalanine metabolism showed no significant changes (Fig. 6F). Furthermore, higher AUC values were observed when grouping the low- and high-BPH hybrids at a more stringent level, while lower values were obtained when dividing the test crosses into approximately equal halves (Supplementary Figs S4-S5). Our results indicated that the 11 identified heterosis-associated pathways, which were significantly enriched at most developmental stages in the diallel crosses, effectively predicted height heterosis in the test crosses.

## Discussion

In this study, we identified robust biomarkers for dynamic height heterosis in rice, based on metabolic pathways significantly enriched at most developmental stages. In alignment with the phenotypic transformations observed in fields, metabolomic changes at the elongation stage contribute more substantially to height heterosis than those at the other stages. Recognizing these critical stages through time-series analysis may facilitate the identification of biomarkers for predicting biomass or height in breeding programs for perennial energy crops and trees (Canas et al., 2015; Maddison et al., 2017).

Among the 11 selected metabolic pathways, all of which were significantly enriched at the elongation stage, pathways related to carbohydrate and amino acid metabolism comprised the majority and exhibited approximately opposite trends in metabolite levels. Given the tight connection between plant height and grain yield in rice (Xue et al., 2008; Chen et al., 2020; Dan et al., 2021), variations in these two types of pathways may determine the optimal balance between vegetative and reproductive traits in the formation of heterosis in hybrid rice (Birchler, 2019; Li et al., 2020; Dan et al., 2021). Furthermore, the strategy of incorporating pathways that are significantly enriched at most developmental stages—rather than exclusively those enriched at all stages—could enhance the robustness of pathway biomarkers for predictions across diverse populations, environments, and developmental stages. We think that this advantage arises from the comprehensive information provided by the predictive features.

While we have identified multiple metabolic pathways associated with dynamic height heterosis, their mechanistic roles require in-depth investigation. Simultaneous mutations of key genes within these metabolic pathways could clarify their functional contributions. Notably, the numbers of heterosis-associated pathways and analytes were greater at the earlier stages than at the later ones, suggesting a more complex and potentially pivotal metabolomic basis for heterosis at early developmental stages. Moreover, our pathway analysis is inherently limited by the coverage of the KEGG database; unannotated pathways may also influence height heterosis. Expanding the catalog of heterosis-associated pathways will refine the metabolomic landscape and improve predictive accuracy.

While the variation patterns of some pathways were consistent across hybrid populations, we observed discrepancies in metabolite levels of certain pathways. The metabolomic heterogeneity between populations (Dan et al., 2021), similar to transcriptional variability among phenotypically similar patients (Menche et al., 2017; Zhou et al., 2019b), may underlie these inconsistencies. Given the observed variations in predictive accuracies, hybrids exhibiting intermediate heterosis warrant further investigation. Comprehensive metabolomics data that incorporate metabolite profiles obtained via both positive and negative ionization modes, as well as through both GC and LC methods, will be essential. The integration of multiple layers of omics information (Weckwerth et al., 2020; Xu et al., 2021), including genomics, transcriptomics, proteomics, metabolomics, and microbiome data, will also optimize pathway biomarkers for predicting complex phenotypes across populations (individuals), environments, and developmental stages.

## Material and methods

### Plant materials

The rice accessions selected for hybridization were previously described (Dan et al., 2020). In brief, two rice hybrid populations were utilized to develop biomarkers for plant height heterosis. The first population adopted a complete diallel cross design involving 18 *indica* and *japonica* accessions as parents, while the second population consisted of 107 test crosses with recombinant inbred lines crossed with the Honglian-type cytoplasmic male sterile line, Yuetai A. The plant height, defined as the distance from the plant base to the highest part of the leaf or panicle, of the diallel cross population was measured with tape measures at four growth stages from seedling to maturation: the seedling stage (28 days after transplantation), elongation stage (47 days after transplantation), flowering stage (time of flowering), and maturation stage (time of maturation). The plant height data for the maturation stage of the diallel cross population were reported in a previous study (Dan et al., 2021), and the complete plant height data are provided in Supplementary table S1. For comparative purpose, several analysis results related to better-parent heterosis for plant height at the maturation stage were retrieved from earlier research (Dan et al., 2021). The plant height of the test cross population was only measured at the maturation stage. The two hybrid populations were grown under different growth conditions and planting densities at the same location (Ezhou, Hubei Province) in 2012 and 2015. Three biological replicates were designed for each genotype, with five plants from the middle region of each row selected for height measurement.

### Metabolite profiling

Untargeted metabolite profiles in the negative ionization mode of 15-d-old parental seedlings were obtained using previously described methods (Dan et al., 2020). In brief, a 1290 Infinity liquid chromatography system (Agilent Technologies), an Agilent quadrupole time-of-flight mass spectrometer (Agilent 6550 iFunnel QTOF), and a Triple TOF 6600 mass spectrometer (AB SCIEX) were employed to acquire the raw metabolomic profiles. The stability of the equipment and the accuracy of the metabolite profiles were assessed using intermittently injected quality control samples. A total of 3,746 analytes were detected, and 114 metabolites were identified through matching with in-house spectral libraries of standards by Shanghai Applied Protein Technology Co., Ltd. To prepare the metabolomic data for statistical analyses, sample normalization (normalization by sum), data transformation (none), and data scaling (auto scaling) were performed using MetaboAnalyst 4.0 (Chong et al., 2018). The hybrid metabolome used for the analyses was calculated by averaging the metabolite levels of the corresponding parents according to previously reported methods (Dan et al., 2019; Dan et al., 2020).

### Identification of heterosis-associated metabolic pathways and analytes

The Metabolite identification and Dysregulated Network Analysis web server (MetDNA version 1.2.1) (Shen et al., 2019), which annotates metabolites based on the rule that neighboring metabolites tend to share similar MS2 spectra, was used to perform pathway enrichment analysis for height heterosis. The hybrids were categorized into high-better-parent heterosis (control group) and low-better-parent heterosis (case group) groups according to quantiles (≥ 75th percentile and ≤ 25th percentile) of the entire hybrid population, and the calculated hybrid metabolome was utilized for dysregulated network analysis. Analysis parameters were set according to the metabolite profiling experiments, with *Arabidopsis thaliana* (thale cress) selected as the species. Student’s *t*-test was employed for univariate statistics, with a cutoff *P*-value of 0.05 (with adjustment). Enriched metabolic pathways and quantitative information on metabolic pathways, which were included in the analysis report, were combined to identify significantly enriched metabolic pathways for height heterosis at each stage. The metabolic pathways that were significantly enriched (*P* values less than 0.05) and exhibited significant differences in metabolite levels between the high-and low-better-parent heterosis hybrids were treated as height heterosis-associated metabolic pathways.

For the identification of heterosis-associated analytes, we employed a partial least square-based strategy (Wold, 1975; Dan et al., 2021). Initially, the number of latent factors in the models was determined based on the highest achieved *r* values. The performance of the predictive models was assessed using 10-fold cross-validation and permutation tests conducted with MetaboAnalyst 4.0 (Chong et al., 2018). Subsequently, average values of variable importance in the projection for all latent factors were calculated to evaluate the contributions of specific analytes to plant height heterosis. Redundant analytes in the predictive models were removed by reducing the numbers of predictive analytes. The final number of predictive analytes for plant height heterosis at the seedling, elongation, flowering, and maturation stages were 300, 300, 100, and 100, respectively.

### Statistical analyses

Better-parent heterosis for plant height was calculated using the equation: BPH = (F_1_-P_H_)/P_H_, where F_1_ represents the plant height of the hybrid and P_H_ denotes the height of the taller parent. The hybrid metabolome was calculated as follows: M_H_ = (M_F_ + M_M_)/2, where M_H_, M_F_, and M_M_ indicate the metabolite levels of the F_1_ hybrid, female parent, and male parent, respectively. Pearson correlation analysis of height heterosis across the four developmental stages was performed using the “Statistical Analysis” module in MetaboAnalyst 4.0 (Chong et al., 2018). Comparisons of height heterosis and metabolite levels of significantly enriched pathways across stages were analyzed using one-way ANOVA (with least significant difference for post hoc multiple comparisons, *P* < 0.05) via SPSS (IBM SPSS Statistics for Windows, Version 20.0. Armonk, NY: IBM Corp.). Analyses comparing plant height heterosis between the diallel and test crosses, as well as metabolite levels between high- and low-BPH hybrids, were conducted using independent samples *t*-test in SPSS (two-tailed). Pearson correlation analysis between height heterosis and the metabolite levels of metabolic pathways was also performed using SPSS. Both univariate and multivariate ROC curve analyses were conducted using the “Biomarker Analysis” module in MetaboAnalyst 4.0 (Chong et al., 2018). The metabolic ratios of the top 100 were computed and included in the subsequent analysis. Random forest analysis (Breiman, 2001) was selected as the classification and feature ranking method for the multivariate ROC curve analysis.

## Author contributions

Z.D. designed the research; Z.D. and W.H. recorded field phenotypic data; Z.D. and Y.C. performed most of the experiments; Z.D. performed the data analyses; W.H. supervised the experiments; Z.D., Y.C., and W.H. wrote the manuscript.

## Data availability

Raw metabolite profiles of experimental and quality control samples and corresponding metabolomic data are available from the MetaboLights database under the accession number MTBLS742.

## Supplementary data

**Supplementary Table S1** Plant height of inbred lines and the F1 hybrids.

**Supplementary Table S2** Metabolite levels of four metabolic pathways for 26 high- and 26 low-BPH test crosses.

**Supplementary Figure S1** Univariate and multivariate ROC curve analysis of different numbers of significantly enriched pathways for height heterosis at seedling stage of the diallel crosses.

**Supplementary Figure S2** Univariate and multivariate ROC curve analysis of the 11 selected metabolic pathways for height heterosis at the three later developmental stages of the diallel crosses.

**Supplementary Figure S3** Multivariate ROC curve analysis of three metabolic pathways for height heterosis of the test crosses.

**Supplementary Figure S4** Validation of pathway biomarkers with the top 15 and bottom 15 test crosses.

**Supplementary Figure S5** Validation of pathway biomarkers with the entire test crosses.

