## Supplementary figures for "Machine learning-based prediction of dynamic height heterosis with pathway biomarkers in rice"

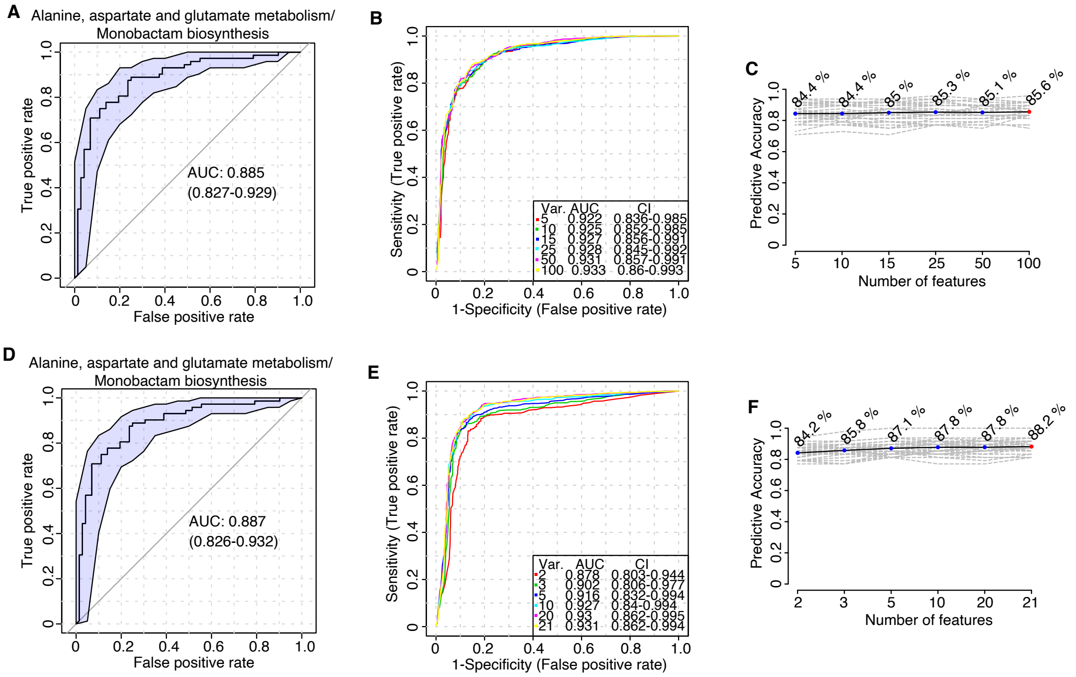


**Fig. S1** Univariate and multivariate ROC curve analysis of different numbers of significantly enriched pathways for height heterosis at seedling stage of the diallel crosses. (**A-C**) Uni- and multi-variate receiver operating characteristic (ROC) curve analysis of the 18 significantly enriched metabolic pathways for height heterosis at the seedling stage. The values of area under the curve (AUC) with a single predictor (A), multiple predictors (B), and changes in predictive accuracies with different numbers of predictive features (C) were shown. (**D-F**) Uni- and multi-variate ROC curve analysis of the six significantly enriched metabolic pathways for height heterosis at all four stages. The values of AUC with a single predictor (D), multiple predictors (E), and changes in predictive accuracies with different numbers of predictive features (F) were shown. VAR., variable. CI, confidence interval.

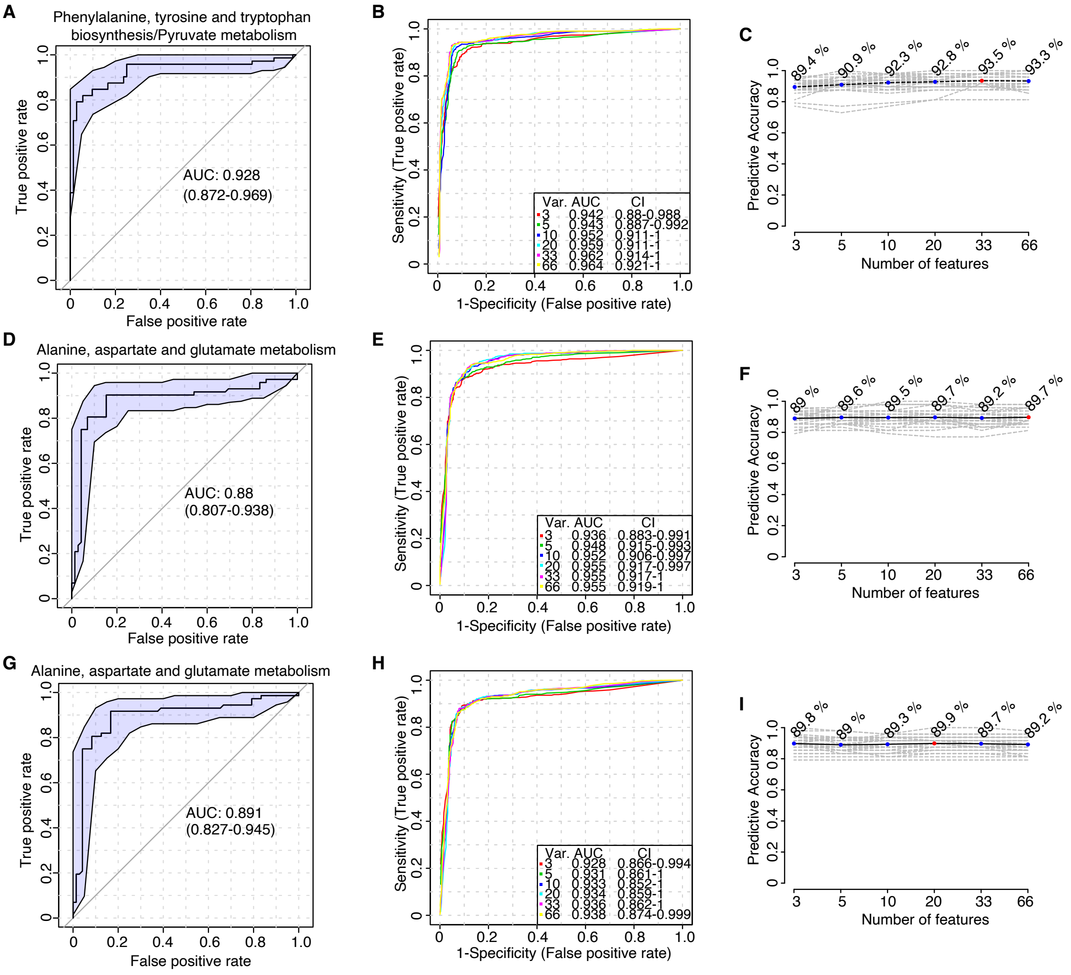


**Fig. S2** Univariate and multivariate ROC curve analysis of the 11 selected metabolic pathways for height heterosis at the three later developmental stages of the diallel crosses. Uni- and multi-variate ROC curve analyses of the 11 selected metabolic pathways for height heterosis at the elongation stage (A-C), flowering stage (D-F), and maturation stage (G-I) were performed. The values of AUC with a single predictor (A, D, and G), multiple predictors (B, E, and H), and changes in predictive accuracies with different numbers of predictive features (C, F, and I) were shown.

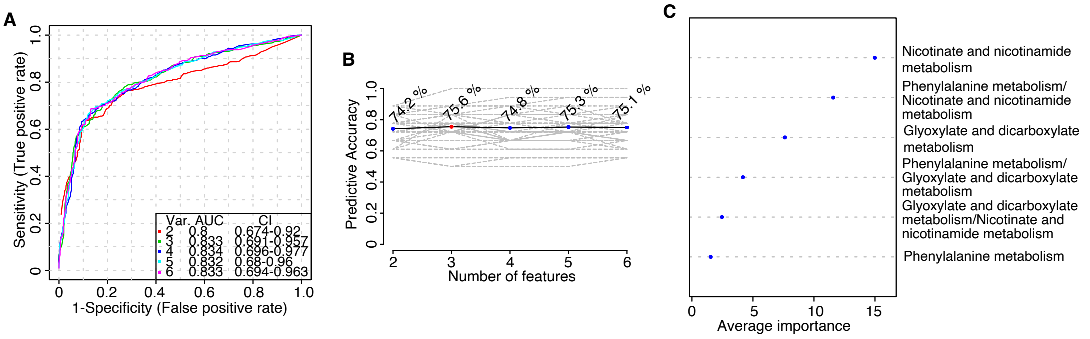


**Fig. S3** Multivariate ROC curve analysis of three metabolic pathways for height heterosis of the test crosses. The test crosses were divided into high- and low-BPH groups based on the quantiles of height heterosis, and the two groups included 26 high- and 26 low-BPH hybrids. Metabolite levels of three pathways that belong to the 11 selected metabolic pathways were obtained through dysregulated network analysis. The values of AUC with multiple predictors (A), changes in predictive accuracies with different numbers of predictive features (B), and average importance of the top six features (C) were shown.

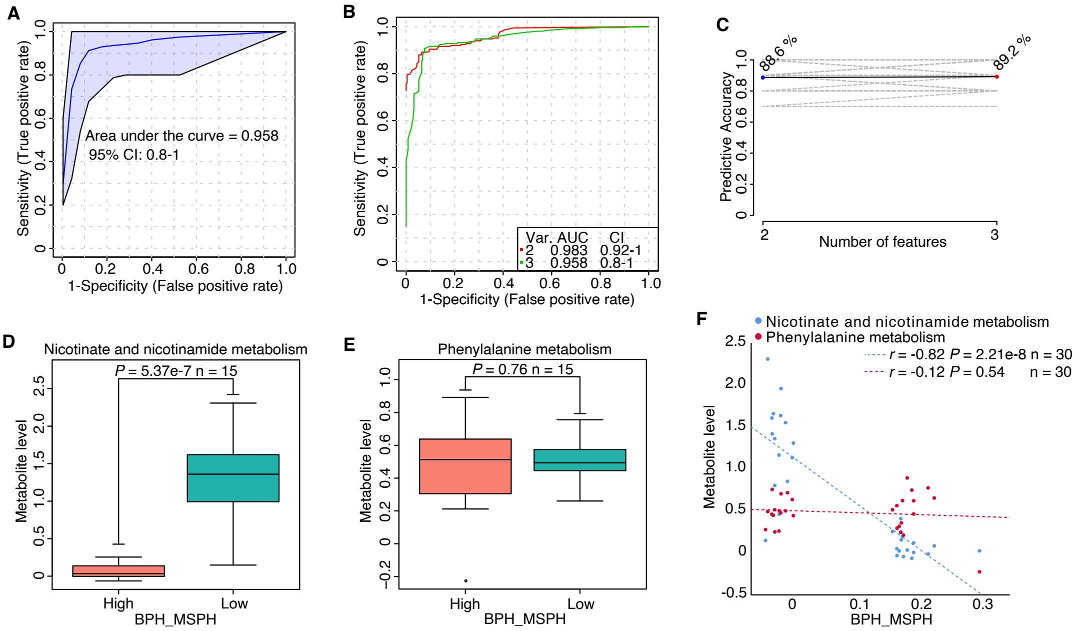


**Fig. S4** Validation of pathway biomarkers with the top 15 and bottom 15 test crosses. (**A-C**) Multivariate ROC curve analysis on the top 15 and bottom 15 test crosses. The top 15 and bottom 15 hybrids from the test crosses were treated as the high- and low-BPH groups. Metabolite levels of two pathways that belong to the 11 selected metabolic pathways were obtained through dysregulated network analysis. The values of AUC with three predictors (A), changes in AUC with different numbers of predictors (B), and predictive accuracies with different numbers of predictive features (C) were shown. (**D-E**) Comparisons of metabolite levels of the two pathways between the high- and low-BPH hybrids. Metabolite levels of nicotinate and nicotinamide metabolism (D) and phenylalanine metabolism (E) between the high- and low-BPH hybrids were compared. (**F**) Correlations between height heterosis of the test crosses and metabolite levels of the two metabolic pathways. For box plots in (D-E), the middle line is the median. The lower and upper boundaries of the boxes are the first and third quantiles, respectively. The whiskers represent the 1.5 times interquartile range and the circles represent the outliers.

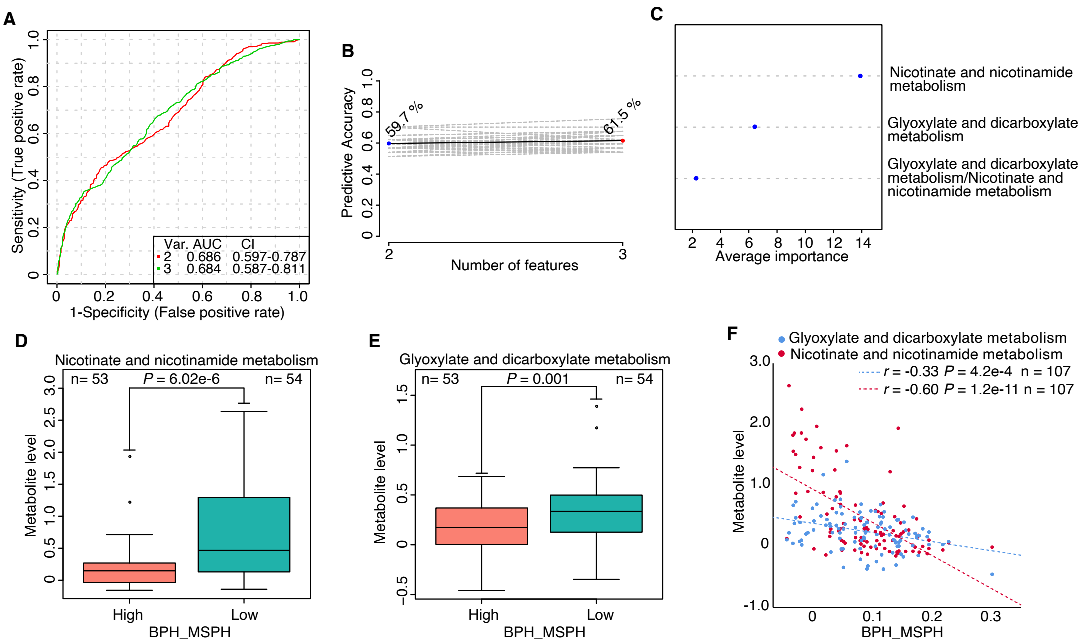


**Fig. S5** Validation of pathway biomarkers with the entire test crosses. (**A-C**) Multivariate ROC curve analysis on all the test crosses. All the test crosses were divided into high- (53) and low- (54) BPH two groups. Metabolite levels of two pathways were obtained through dysregulated network analysis. The values of AUC with multiple predictors (A), changes in predictive accuracies with different numbers of predictive features (B), and average importance of the top three features (C) were shown. (**D-E**) Comparisons of metabolite levels of the two pathways between the high- and low-BPH hybrids. Metabolite levels of nicotinate and nicotinamide metabolism (D) and glyoxylate and dicarboxylate metabolism (E) between the high- and low-BPH hybrids were compared. (**F**) Correlations between height heterosis of the test crosses and metabolite levels of the two metabolic pathways. For box plots in (D-E), the middle line is the median. The lower and upper boundaries of the boxes are the first and third quantiles, respectively. The whiskers represent the 1.5 times interquartile range and the circles represent the outliers.
